# 3D super-resolution deep-tissue imaging in living mice

**DOI:** 10.1101/790212

**Authors:** Mary Grace M. Velasco, Mengyang Zhang, Jacopo Antonello, Peng Yuan, Edward S. Allgeyer, Dennis May, Ons M’Saad, Phylicia Kidd, Andrew E. S. Barentine, Valentina Greco, Jaime Grutzendler, Martin J. Booth, Joerg Bewersdorf

**Affiliations:** Department of Biomedical Engineering, Yale School of Engineering & Applied Science, CT 06520, USA; Department of Cell Biology, Yale School of Medicine, CT 06520, USA; Interdepartmental Neuroscience Program, Yale School of Medicine, CT 06520, USA; Department of Neuroscience, Yale School of Medicine, CT 06520, USA; Department of Engineering Science, University of Oxford, Oxford OX1 3PJ, UK; Department of Neurology, Yale School of Medicine, CT 06520, USA; Department of Genetics, Yale School of Medicine, CT 06520, USA; Department of Dermatology, Yale Stem Cell Center, Yale Cancer Center, Yale School of Medicine, CT 06520, USA

## Abstract

Stimulated emission depletion (STED) microscopy enables the three-dimensional (3D) visualization of dynamic nanoscale structures in living cells, offering unique insights into their organization. However, 3D-STED imaging deep inside biological tissue is obstructed by optical aberrations and light scattering. We present a STED system that overcomes these challenges. Through the combination of 2-photon excitation, adaptive optics, far-red emitting organic dyes, and a long-working distance water-immersion objective lens, our system achieves aberration-corrected 3D super-resolution imaging, which we demonstrate 164 µm deep in fixed mouse brain tissue and 76 µm deep in the brain of a living mouse.

## Introduction

Much like the invention of the light microscope itself, the advent of super-resolution fluorescence microscopy has triggered paradigm shifts in the way we study biology [1]. In an era where imaging resolution is no longer dictated by Abbe’s law of diffraction [2], the noninvasive visualization of nanoscale dynamic structures in living cells is now attainable [3]. Of the super-resolution imaging modalities, stimulated emission depletion (STED) microscopy [4] is most readily extended to imaging thick tissue samples. The confocal pinhole integrated into most STED microscopes, provides an optical sectioning effect that has enabled super-resolution imaging in fruit flies [5], whole worms [6], and even living mice [7].

The typically ring-shaped depletion focus of a STED microscope is most often generated by applying a 0 to 2π azimuthal phase ramp (or “vortex”) to the depletion beam [8] using either a static phase plate [9] or, more recently, a spatial light modulator (SLM) [10]. Unfortunately, the depletion effects of this focus are limited to the lateral (*xy*) direction, such that the effective point spread function (PSF) remains diffraction-limited in the axial (*z*) direction.

To improve the axial resolution of STED microscopy, two methods have proven useful. First, the STED principle can be combined with a 4Pi geometry featuring two opposing objective lenses [11]. 4Pi-STED systems have been able to achieve three-dimensional (3D) isotropic ∼30 nm super-resolution [12], earning them the name “isoSTED” [13]. However, isoSTED microscopes, in addition to being optically complex, are also typically limited to thin samples. Moreover, the need for opposing objectives precludes their application in larger specimens such as living mice.

An alternative approach to 3D-STED microscopy is the use of a single-objective geometry with a radially symmetric “top-hat” depletion phase mask, so named for the central π-step [14]. The top-hat phase mask generates a depletion focus with two high-intensity lobes above and below the focal plane for fluorescence depletion along the optical axis. While the axial resolution achieved in this way does not match that of an isoSTED microscope, the single-objective geometry permits the imaging of thick samples, including living mice. Furthermore, the top-hat phase mask can be combined with a traditional vortex phase mask to overcome the weak lateral depletion achieved with the top-hat focus alone [15].

However, 3D-STED microscopes that exploit the top-hat phase mask still find limited application in deep-tissue imaging experiments. This is primarily due to the susceptibility of the top-hat depletion focus to optical aberrations [10, 16], which arise from refractive index mismatches between the objective lens immersion medium and the specimen. If left uncorrected, aberrations can raise the intensity minimum in the center of the depletion focus, causing fluorescence emission to be depleted entirely, rather than being confined to the center of the depletion ring. Recently, Heine *et al.* replaced the oil or glycerol immersion objective lens of a typical 3D-STED microscope with a water immersion objective to minimize the spherical aberration induced when imaging aqueous living specimens [17]. Despite the lower NA of water immersion objectives, the authors resolved 153 nm structures axially. Nevertheless, the imaging depths achieved were still limited by scattering, the objective working distance and specimen-induced aberrations (in addition to spherical aberration) that remained uncorrected.

To address scattering, 3D-STED microscopy can be combined with two-photon excitation (2PE) [18, 19], which utilizes near-infrared (NIR) excitation wavelengths with lower scattering cross-sections. Moreover, recent developments in STED microscopy have exploited robust far-red emitting organic dyes which, in addition to being excitable via 2PE [20], also require NIR depletion wavelengths. The shift of both the excitation and depletion light to the NIR regime not only minimizes scattering but also alleviates sample photodamage [21], making these wavelengths better suited for imaging live specimens.

Nevertheless, the presence of optical aberrations is still a major concern for deep-tissue 3D-STED microscopy, even when it is coupled with 2PE. Biological tissue can be very optically heterogeneous and the refractive index is typically highly varying in space. While clearing the sample can address this issue [22], clearing methods are not compatible with living specimens. A gentler approach that is non-destructive to the sample is the use of adaptive optics (AO). In AO, a corrective element such as an SLM or a deformable mirror (DM) is imaged onto the back focal plane of the objective lens and is programmed to impart a phase variation that is equal but opposite to that induced by the sample. The cumulative phase at the sample plane is thus zero and aberration-free excitation and depletion foci can be recovered. This active approach to aberration-correction is ideal for imaging biological specimens where complex, highly varying refractive index maps are the norm [23, 24].

Previous implementations of AO in 3D-STED microscopy have determined the corrective phase modulation using a metric-based approach [10, 25, 26]. The applied correction is systematically adjusted to optimize the quality of the final image. For a given aberration mode, a sequence of images is acquired, with each image corresponding to a different mode coefficient. Each image is then quantified using an image quality metric and the optimal amount of correction is estimated as that which maximizes this metric. This approach, when combined with 3D-STED microscopy, has enabled aberration-corrected imaging of the complete 15 µm thick mitotic spindle in fixed cells [26] and of fluorescent beads through 25 µm of fixed zebrafish retina tissue [10].

Alternatively, wavefront sensing (WFS) can be used to directly measure the sample-induced aberrations. Conveniently, in 2PE microscopes, the guide star required for this approach is provided by the two-photon-excited fluorescent volume [27, 28], which is inherently confined in three dimensions. Fluorescence from this “non-linear guide star” is descanned and directed to a Shack-Hartman sensor (SHS). The SHS then samples the wavefront using a microlens array and generates a spot diagram on a camera positioned at the focal plane of the microlenses. If the wavefront is aberrated, the spots will be displaced from the center of their designated sub-regions. Using these displacements, the SHS can reconstruct the wavefront, determine its constituent Zernike modes and thus the corrective phase that needs to be added to the system. This approach to aberration correction has been used to recover the optimal resolution of 2PE laser scanning [27, 28] and structured illumination [29, 30] microscopes deep in tissue, including in a living mouse [30, 31].

Here, we present a 2PE-STED microscope capable of 3D sub-diffraction-limit resolution deep in aberrating tissue. Its capabilities are made possible by the combined effect of 2PE, far-red emitting organic dyes, WFS-based aberration-correction, and a long-working-distance water immersion objective lens. We demonstrate the system’s capabilities by visualizing the 3D chromatin structure of keratinocytes in fixed mouse skin tissue. We then demonstrate *in vivo* 3D-STED imaging in living mice, using a labelling strategy that enables neuronal labeling of the intact mouse brain with ATTO590, a photostable, live-cell and STED-compatible organic dye [32].

## Results

### Optical Setup

A schematic of our AO- and 2PE-enabled 3D-STED microscope is presented in **Fig. 1(a)** (see details in **Methods**). Our setup is built around a custom upright microscope stand and features a 25×, 1.05 NA water immersion objective lens with a 2 mm working distance. For depletion, we use an 80 MHz repetition rate, 775 nm pulsed laser with a pulse length of ∼600 ps. To impart both the vortex and top-hat phase masks on the same depletion beam, we adopt a double-pass SLM configuration as described by Lenz *et al.* [15]. We also apply a blazed grating pattern on the SLM, to isolate and block any unmodulated light.

**Fig. 1.**
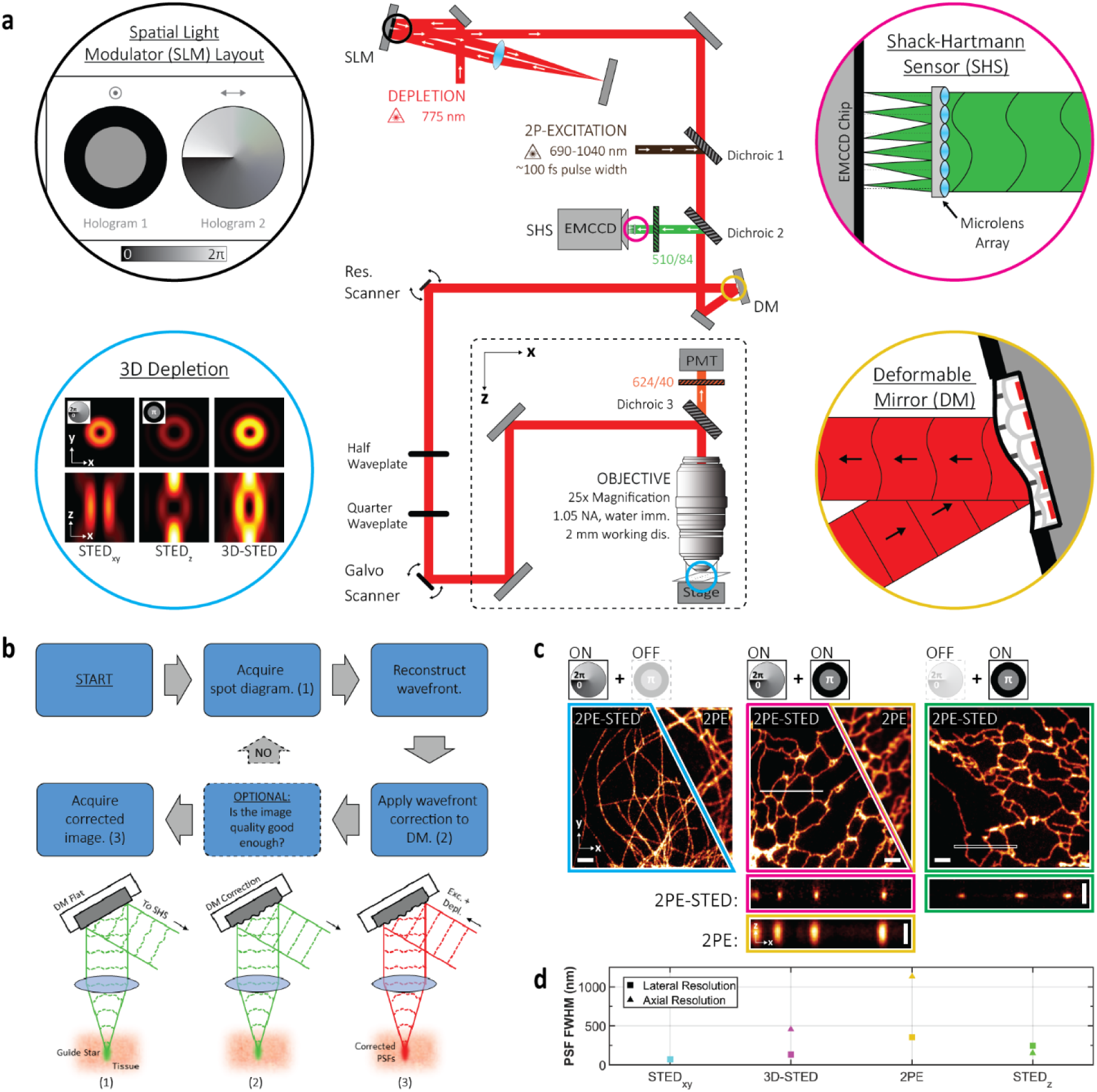
Principles of aberration-corrected 3D-2PE-STED microscopy. **(a)** Simplified schematic of the optical setup. Detailed schematic is available in Fig. 1-figure supplement 1. **(b)** Aberration-correction routine. **(c)** 2PE and 2PE-STED images of ATTO594-labelled microtubules (left) and ER tubules (middle and right) in fixed COS7 cells. Depletion via either or both the vortex and top-hat phase masks was used, as indicated by the schematic above each image. A 3D Gaussian blur of σ_xy_ = 39 nm and σ_z_ = 50 nm was applied to all images. *xy* images depict either a single frame (cyan) or a maximum intensity projection (MIP) of an image stack (magenta and green). *xz* images depict MIPs of the regions demarcated by the white box in the corresponding *xy* image, resliced in the *xz* direction. Scale bars are 2 µm in the *xy* images and 1 µm in the *xz* images. **(d)** Plots summarizing the lateral and axial resolution of the system. See main text and methods for details. Data points are colored to match the border color of the corresponding image in (c).

For 2PE, we use light from a fs-pulsed titanium-sapphire laser that is merged with the STED beam via a dichroic mirror. The two beams are then raster-scanned across the sample by a 10 kHz resonance mirror synchronized with a galvanometric mirror. Fluorescence is collected by the same objective lens that focuses the excitation and depletion light into the sample. Far-red fluorescence emission wavelengths are detected in a non-descanned configuration by a photomultiplier tube (PMT), while green fluorescence emission wavelengths are descanned and directed to a SHS.

Two design features are particularly beneficial for imaging in aberrating tissue. First, we employ a water immersion objective lens that is more closely index-matched to living tissue than oil immersion objective lenses, thus reducing specimen-induced aberrations. Furthermore, the long 2 mm working-distance of this objective lens permits long penetration depths into the specimen and accommodates a wide range of specimen dimensions and configurations. Second, we adopt an AO architecture based on WFS to correct for specimen-induced aberrations that cannot be addressed by simply matching the objective lens immersion medium to the sample. Following the developments of Wang *et al.* [27, 28, 31], we rely on 2PE to generate the guide star required for this WFS approach. Our correction routine is outlined in **Fig. 1(b)**. First, GFP or fluorescein is excited via 2PE to generate a guide star in the sample and the fluorescence from the guide star is descanned to a SHS to generate a spot diagram. The spot diagram is then analyzed to reconstruct the aberrated wavefront. The opposite of the measured aberration modes is then added to the common beam path for aberration correction using a DM positioned in a plane conjugate to the back pupil of the objective lens. This step recovers (close to) aberration-free excitation and depletion foci at the sample such that a super-resolution image can be acquired in the far-red detection channel. In contrast to the approach by Wang *et al.* and inspired by the more recent implementation by Zheng *et al.* [29], the DM and SHS in our system are arranged in a closed feedback-loop. This enables measurement of any residual aberrations after correction and allows the user to optionally validate the correction and implement further rounds of correction, if necessary. This guarantees a more robust and accurate aberration correction step.

To benchmark the resolution capabilities of our system, we imaged fixed COS7 cells featuring ATTO594-labelled microtubules or endoplasmic reticulum (ER) tubules (**Fig. 1(c)**). Intensity profiles extracted from the raw tubule images were then fit using nested-loop ensemble PSF (NEP) fitting [33] unless the tubule could be assumed to be much smaller than the PSF (see **Methods**).

Due to the double-pass configuration of the SLM and the tunable allocation of the depletion laser power between the vortex and top-hat phase masks, lateral and axial resolution are inversely correlated in our system, i.e. allocating more laser power to the vortex phase mask to improve the lateral resolution compromises the axial resolution and *vice versa*. The maximum lateral resolution achievable by the system, obtained by allocating the laser power entirely to the vortex phase mask, was quantified to be 70 nm (**Fig. 1(c)** left; see **Methods**). The maximum axial resolution achievable by the system using only the depletion beam component encoded with the top-hat phase mask, was quantified to be 151 nm (**Fig. 1(c)** right; see **Methods**). These values are 4.2 and 6.5 times below the theoretical lateral and axial diffraction-limited resolution of our 2PE microscope, respectively (296 nm and 988 nm; NA = 1.05, refractive index = 1.33, wavelength = 810 nm) [34].

For 3D-STED imaging, it is expected that the depletion power will be distributed between the vortex and top-hat phase masks according to the resolution requirements of the given experiment. Therefore, 3D-STED resolution (in non-aberrating samples) will be a compromise between the 70 nm and 142 nm lateral and axial values described above (e.g. **Fig. 1(c)** middle).

### 3D-STED Microscopy of Chromatin in Mouse Skin Tissue

To demonstrate the aberration-correction capabilities of our system, we imaged epithelial nuclei in fixed mouse skin tissue. The tissue was harvested from transgenic mice expressing histone 2B-green fluorescent protein (H2B-GFP) under the control of the keratin 14 (K14) promoter. The H2B-GFP fusion proteins were then immuno-labelled with anti-GFP nanobodies conjugated to ATTO594. While the GFP fluorescence was used to generate the non-linear guide star for aberration correction, the ATTO594 fluorescence enabled the 3D super-resolution imaging of chromatin within the epithelial nuclei.

In **Fig. 2(a)**, we compare corrected and uncorrected (for specimen-induced aberrations) 3D-STED datasets acquired from the same volume, 62 µm deep in the labelled tissue. Instrument aberrations were corrected for in both cases. For STED imaging, only the top-hat phase profile was used for depletion because the size scale of the labelled chromatin structures was within the diffraction-limited lateral but not axial resolution of conventional 2PE microscopy. Since the top-hat depletion focus is more sensitive to aberrations than the vortex depletion focus, this imaging configuration was still sufficient for demonstrating the benefits of our AO approach. Indeed, the finer chromatin structures, indistinguishable without AO (**Fig. 2(a)**, left-top), were clearly resolved when aberration-correction was applied (**Fig. 2(a)**, left-bottom). This improvement is even more striking in the axial direction. We compare the uncorrected and corrected *yz* images (**Fig. 2(a)**, right) extracted from the area marked by the white box (**Fig. 2(a)**, left). The corrected image clearly shows details in the chromatin structure that are lost in the uncorrected image, and due to the improved axial resolution granted by the STED effect, the details no longer appear elongated as they do in the 2PE image. These effects are also illustrated by the intensity profiles in **Fig. 2(d)** of the areas marked by the dashed lines in **Fig. 2(a)**. The plots show that the fluorescent signal is almost entirely absent in the uncorrected image, likely depleted by the compromised STED focus, but is recovered in the corrected image. This improvement cannot be attributed to photobleaching as the uncorrected image was acquired first.

**Fig. 2.**
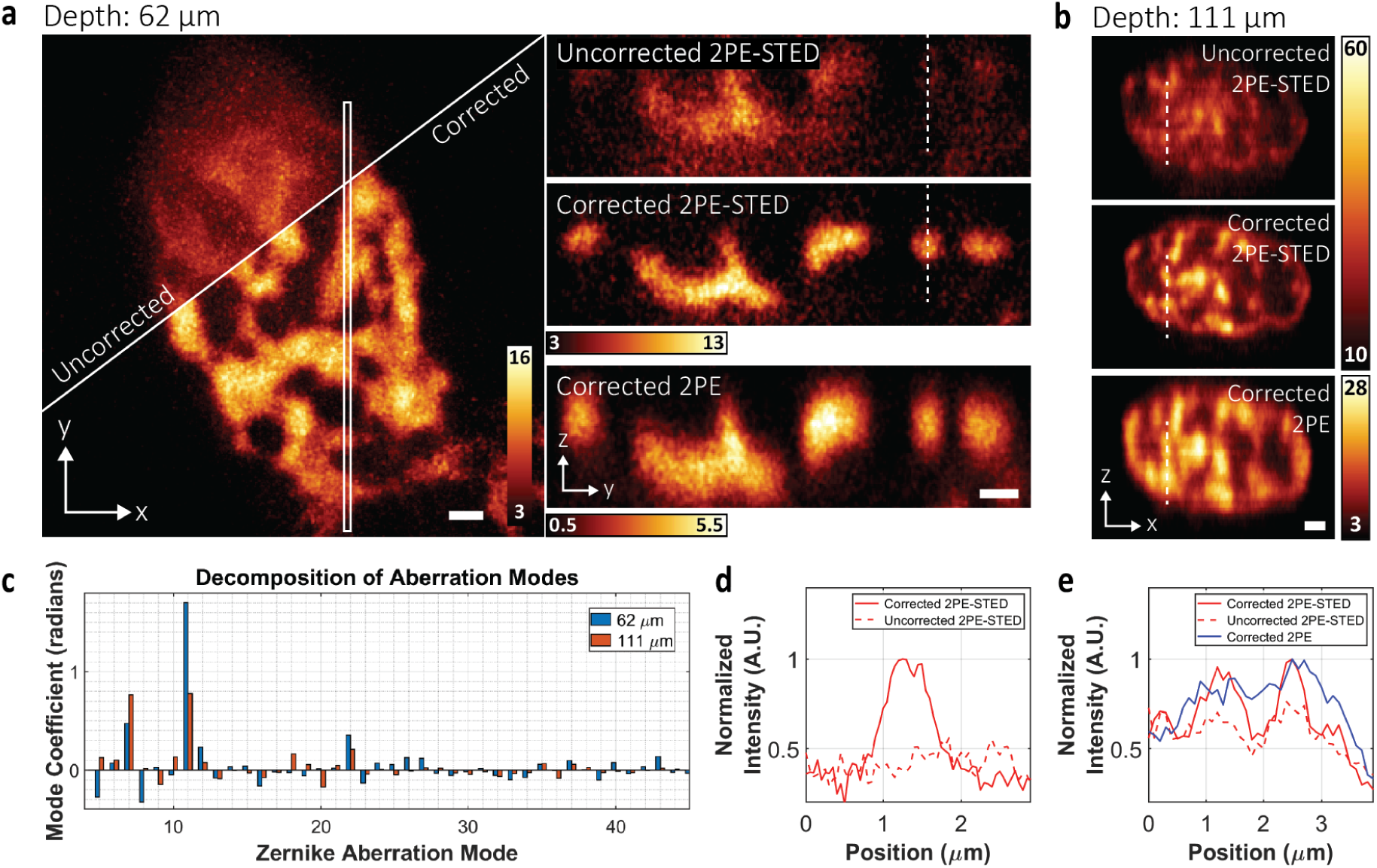
Aberration-corrected 2PE-STED microscopy in mouse skin tissue. **(a)** H2B-GFP distribution within the nucleus of an epithelial cell located 62 µm below the tissue surface. **Left:** MIP of the uncorrected (top) and corrected 2PE-STED (bottom) image stacks. **Right:** MIP of the region in the white box, resliced in the y*z* direction from the uncorrected 2PE-STED (top), corrected 2PE-STED (middle), and corrected 2PE (bottom) image stacks. **(b)** MIP of uncorrected 2PE-STED (top), corrected 2PE-STED (middle), and corrected 2PE (bottom) image stacks, resliced in the *xz* direction. Cell was located 111 µm below the tissue surface. In both **(a)** and **(b)**, depletion via only the top-hat phase mask was used. All images were smoothed using a 3D Gaussian blur of σ_xy_ = 39 nm and σ_z_ = 50 nm for (a), and 100 nm for (b). Scale bars: 1 µm. **(c)** Zernike mode decomposition (modes 5 to 45) of the DM correction applied for acquiring the corrected image stacks in (a) (blue) and (b) (orange). Radians are defined with respect to λ = 520 nm. **(d)** Plot of the intensity profile at the positions marked by the white dashed lines in (a). The solid red line corresponds to the corrected 2PE-STED image. The dashed line corresponds to the uncorrected 2PE-STED image. **(e)** Plot of the intensity profile at the positions marked by the white dashed lines in (b). The solid red line corresponds to the corrected 2PE-STED image. The dashed red line corresponds to the uncorrected 2PE-STED image. The solid blue line corresponds to the corrected 2PE image. For (d) and (e), the profiles were acquired from a sum intensity projection of the raw, unsmoothed data.

To induce more aberrations and scattering for test purposes, a thicker skin tissue section was prepared and this time was mounted upside-down so that visualization of the epithelial chromatin required aberration-corrected imaging through the adipose tissue underlying the epidermis. In this sample, our imaging volume was located 111 μm below the surface of the tissue section. Nevertheless, our AO approach still improved the aberrated image quality (**Fig. 2(b)**, top) to a level where the chromatin structure could be clearly resolved (**Fig. 2(b)**, middle). Furthermore, the axial resolution was sufficient to discern details that were otherwise blurred together in the diffraction-limited 2PE image (**Fig. 2(b)**, bottom). These observations are confirmed by line profile intensity plots (**Fig. 2(e)**) measured from the areas marked by the dashed lines in **Fig. 2(b)**.

For our AO approach, it is imperative that the acquired SHS spot diagram has a sufficient signal-to-noise level so that reliable wavefront reconstruction can be performed (**Fig. 2(c)**; following the Zernike numbering convention of Noll [35]). In the non-inverted tissue sections, the GFP signal was sufficiently bright such that spot diagrams could be acquired in as little as 1.5 s. In the inverted skin tissue samples, we found that light scattering, likely from the adipose tissue, lowered the signal level of the spot diagrams. We therefore increased the excitation power from 3.02 to 11.29 mW and the SHS camera exposure time to 12 s to increase the detected signal.

### 3D-STED Microscopy of Astrocytes in Mouse Brain Tissue

To demonstrate our system’s capabilities in another aberrating sample, we imaged astrocytes in a 300 µm thick mouse brain tissue section. Glial fibrillary acidic proteins (GFAPs), markers for astrocytes, were immuno-stained with ATTO594. The labelled tissue was then mounted in fluorescein, so that a guide star could be generated from the green fluorescence using 2PE. For STED imaging, only the top-hat depletion profile was applied for axial resolution enhancement.

In **Fig. 3(a)**, uncorrected and corrected 2PE-STED and corrected 2PE *xz* frames are shown from an imaging volume located 164 μm below the tissue surface (see also **Supplementary Movie 1**). Aberrations had a detrimental effect on the intensity and resolution of the uncorrected image. However, aberration correction (**Fig. 3(d)**) improved this such that the average intensity within the cyan and magenta boxes increased by a factor of ∼8.8 (**Fig. 3(b)**). This effect is also exhibited in the line profile intensity plots (**Fig. 3(c)**). The plots also show that two astrocyte branches that appear as one in the 2PE image are distinguishable in the corrected 2PE-STED image, elucidating the STED effect.

**Fig. 3.**
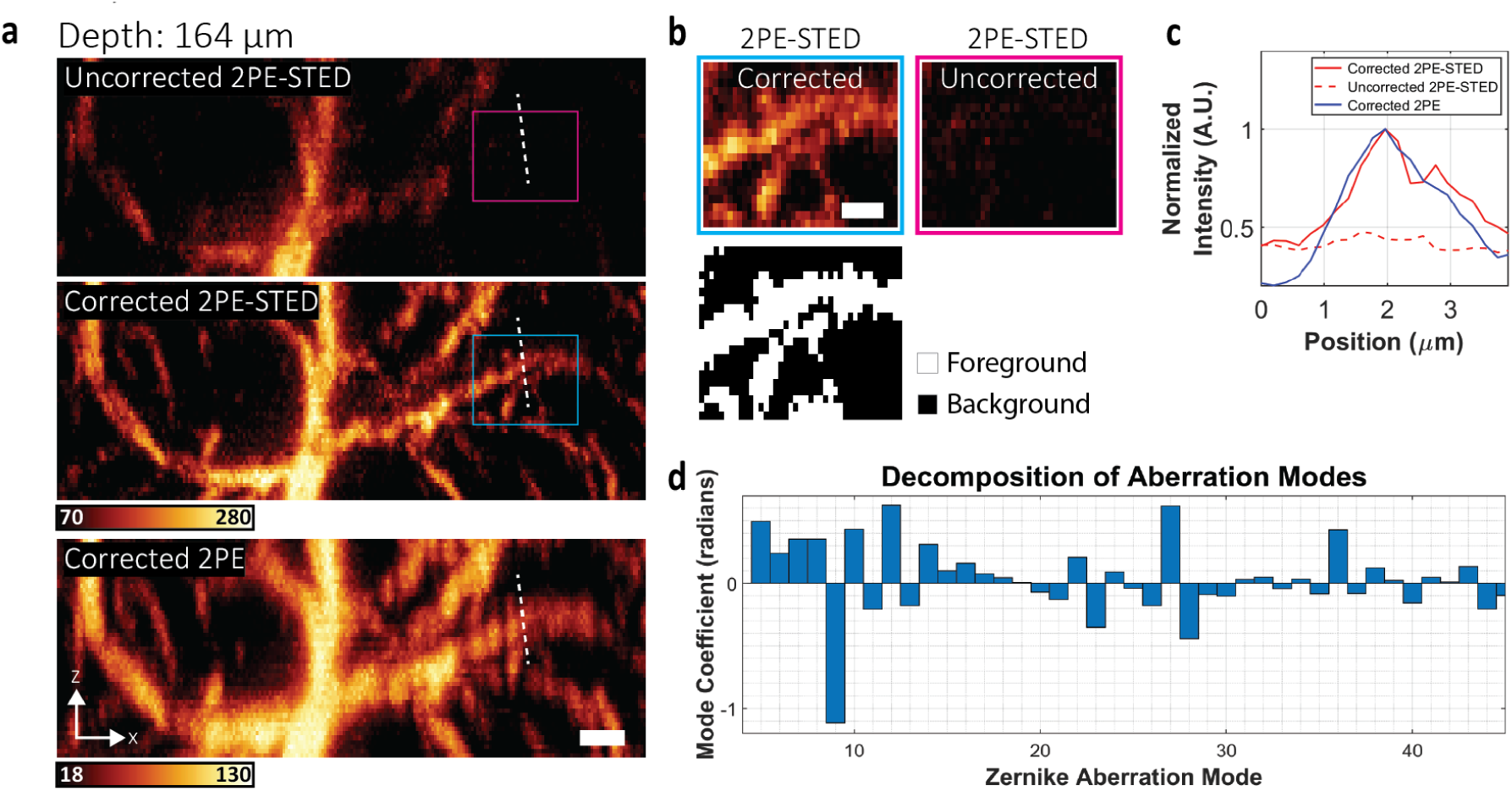
Aberration-corrected 2PE-STED microscopy in fixed mouse brain tissue. **(a)** MIP of uncorrected 2PE-STED (top), corrected 2PE-STED (middle), and corrected 2PE (bottom) image stacks (unsmoothed) of ATTO594-labelled astrocytes, resliced in the *xz* direction. The center of the image stack was 164 µm below the tissue surface. Scale bar: 2 µm. **(b) Top:** Areas corresponding to the magenta and cyan boxes in (a) showing an ∼8.8-fold increase in foreground intensity between the uncorrected and corrected images. **Bottom:** Mask used to delineate the foreground and background pixels. The mask was generated using the Otsu method. The intensity of each image was calculated as the mean of the foreground pixels minus the mean of the background pixels. Scale bar: 1 µm **(c)** Plot of the intensity profiles at the positions marked by the white dashed lines in (a). The solid red line corresponds to the corrected 2PE-STED image. The dashed line corresponds to the uncorrected 2PE-STED image. The solid blue line corresponds to the corrected 2PE image. The profiles were acquired from a sum intensity projection of the raw, unsmoothed data. **(d)** Zernike mode decomposition of the DM correction applied for acquiring the corrected image stacks in (a).

### Aberration-corrected 2PE-STED imaging of neurons in a living mouse

To demonstrate the full potential of our super-resolution deep-tissue imaging system, we performed aberration-corrected, 3D-2PE-STED imaging in the intact brain of a living mouse. For these experiments, we required an *in vivo* labelling procedure that targeted neurons in the brain with the far-red live-cell compatible dye ATTO590. It has previously been shown that labelling with organic dyes emitting in the red to far-red range is advantageous for *in vivo* STED microscopy, as these dyes exhibit superior photophysical properties compared to their red fluorescent protein counterparts [36]. In line with this finding, we used a wild-type CD1 mouse line and recombinant adeno-associated virus (rAAV, serotype 2) infection to induce the expression of fused cytosolic GFP and Halo-tags [37] in a subset of cortical neurons (**Fig. 4(a)**). While the GFP provided the fluorescence for guide star generation, the expressed Halo-tags were labelled with Halo-reactive ATTO590-chloroalkane (ATTO590-CA) for 2PE-STED imaging (**Fig. 4(b)**).

**Fig. 4.**
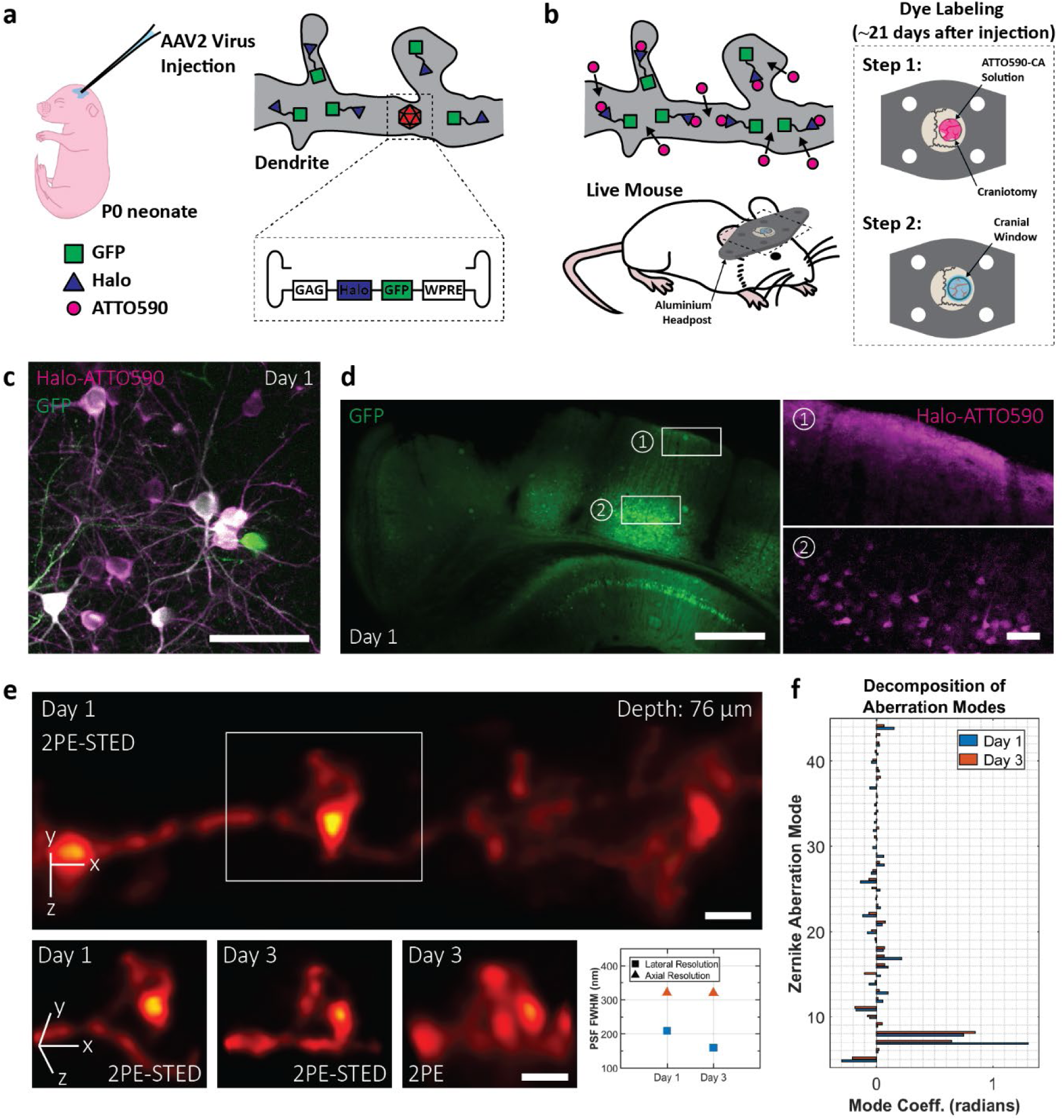
2PE-STED imaging of dendritic spines *in vivo*. **(a)** and **(b)** Strategy for neuron labelling in a living mouse using ATTO590. **(c)** *In vivo* 2PE overview image acquired on a commercial 2PE system 1 day after labelling. Green and magenta correspond to GFP and ATTO590 signal, respectively. Scale bar: 50 μm. **(d)** Widefield images demonstrating the spatial extent of the rAAV infection (represented by GFP labelling; left), and ATTO590 labelling (right). ATTO590 images correspond to the regions outlined by white boxes in the GFP image. Scale bars: 500 μm (left) and 50 μm (right). **(e) Top:** Aberration-corrected STED image stack of a dendrite 76 μm below the cortical surface. Depletion via both the vortex and depletion phase masks was used. A 3D Gaussian blur of σ_xy_ = 60 nm and σ_z_ = 75 nm was applied. Scale bar: 1 μm. **Bottom:** Repeated imaging of the dendritic spine highlighted by the white box in (e). Aberration-corrected imaging was performed 1 (first panel) and 3 (second and third panels) days after the labelling. Scale bars: 1 μm. Resolution values, as quantified using NEP fitting, are shown in the fourth panel. Blue and orange markers correspond to days 1 and 3, respectively. On day 1, lateral and axial PSF FWHMs were 209 and 321 nm. On day 3, they were 160 and 320 nm. **(f)** Zernike mode decomposition of the DM correction applied for acquiring the image stacks in (e). Blue and orange bars correspond to days 1 and 3, respectively.

Two-color imaging of the brain surface with a conventional 2PE microscope, one day after labelling, revealed bright neurons in both the green and red channels (**Fig. 4(c)**). Despite the non-fluorogenic nature of ATTO590, 2PE imaging with excellent signal-to-noise was possible as deep as 174 μm below the cortical surface, confirming the compatibility of our labelling procedure with the imaging depths accessed by our system.

To confirm the spatial extent of neuronal labelling, we performed wide-field imaging of fixed coronal brain sections, harvested 1 day after labelling (**Fig. 4(d)**). rAAV infection, indicated by the GFP signal (**Fig. 4(d)**, left), was evident throughout the injected hemisphere, and the topical application of ATTO590-CA was sufficient to label neuronal cell bodies down to ∼650 µm below the brain surface (**Fig. 4(d)**, right).

Next, we imaged ATTO590-labelled neurons in anesthetized mice with our 3D-2PE-STED instrument. In **Fig. 4(e)**, we show an aberration-corrected 3D-2PE-STED image stack of a dendrite located 76 μm below the cortical surface (see also **Supplementary Movie 2**). The 3D image volume is displayed at a 70° rotation about the *x*-axis to emphasize the 3D resolution achieved. Both the vortex and top-hat depletion phase masks were used to acquire this dataset. The white box (**Fig. 4(e)**, top) highlights a dendritic spine that bends in and out of the *xy*-plane, highlighting the need for 3D imaging to capture its complete morphology (**Fig. 4(e)**, bottom, first panel). The same spine was re-imaged two days later (**Fig. 4(e)**, bottom, second panel), and it exhibited morphological changes in the region of the spine head, which is possibly reconnecting with the main dendritic branch. Such details are lost when using (aberration-corrected) 2PE microscopy (**Fig. 4(e)**, bottom, third panel) due to the inferior resolution, especially in the axial direction. Resolution quantifications are summarized in **Fig. 4(e)** (bottom, fourth panel).

For a typical experiment, the image volume was recorded as a set of 20 μm × 20 μm frames (512 × 512 pixels and 39 nm pixel size) spaced 50 nm apart axially. Each frame took 10 s to acquire. For some areas, “jittering” artifacts could be observed that were a result of the animal’s breathing and heartbeat. If the motion was not too severe, it could be offset by offline image registration. In this case, each step in the image stack was acquired as a set of 10 individual frames, each acquired in 1 s. Each frame set was registered and averaged before being assembled into the final image stack.

However, it is preferable to mitigate the motion itself. This was achieved by positioning the coverslip sealing the craniotomy in direct contact with the brain surface, thereby suppressing tissue motion. Moreover, we avoided imaging regions near major arteries as the blood flowing through the arteries can agitate the surrounding tissue. By observing these guidelines, motion artifacts could be sufficiently diminished such that offline image registration was not required.

In **Fig. 4(f)**, we show the mode decomposition of the wavefront correction applied on the DM. The major corrected aberration was coma, which likely stemmed from a tilt of the cranial window. It has previously been shown that coverslip tilt can have deleterious effects on image quality by compromising the static spherical aberration correction applied via the objective correction collar [38, 39]. Since it is very difficult to control, and near-impossible to eliminate window tilt during animal surgery and mounting, having a fast and adaptive means of aberration-correction to compensate for this effect is crucial for 3D STED *in vivo* imaging.

### Summary and Outlook

Here, we present a microscope that delivers sub-diffraction-limit imaging resolution, in three dimensions, deep in biological tissue. We demonstrate these capabilities 164 μm deep in fixed mouse brain tissue and 76 μm deep in the brain of a living mouse. Our imaging depths were practically limited by light scattering of the fluorescence signal that was used for the SHS wavefront measurements. Further depth-improvements can be envisioned by using sensorless aberration correction approaches alone or by shifting the SHS wavefront detection wavelength into the red emission range [31]. Additional studies will also be needed to systematically determine the achievable depth-dependent resolution in different tissues.

For our *in vivo* imaging experiments, we combined the use of organic dyes, Halo-tags, and rAAV technology to label neurons deep in the living mouse brain. While live-cell compatible red organic dyes like ATTO590 offer superior STED resolution and photostability over fluorescent proteins of the same color [36], the use of rAAVs offers flexibility in the labelling scheme. The virus can be easily modified to target a different cell type, to label specific proteins (e.g. by expressing the Halo-tags fused to a protein of interest), and to additionally express SNAP-tags so that a second live-cell compatible dye (e.g. silicon rhodamine [36, 40]) can be used for multicolor super-resolution imaging. To simplify future experiments, transgenic mouse lines that express SNAP- and/or Halo-tags constitutively [36] offer a user-friendly alternative. Moreover, instead of topically applying the cell-permeable dye to the exposed brain surface, one may consider intravenously injecting dyes that are able to cross the blood-brain barrier [41]. Either or both modifications to the labelling protocol will minimize the exposure and manipulation of the imaged tissue.

It is encouraging that in our *in vivo* STED experiments we did not observe any severe structural changes in the imaged neurons that could be linked to phototoxicity. This may exemplify the live-cell imaging benefits of using a resonant scanner and red-shifted excitation and depletion wavelengths [21]. Nevertheless, the potential negative effects of labeling and imaging on the physiology of the mouse brain will require further detailed investigation.

The possible applications of our microscope are extensive. For example, it enables the study of neuronal plasticity in deeper layers of the brain cortex, the dynamic nanoscale organization of complex structures such as glomeruli in kidney or the structural reorganization of chromatin during cell differentiation in tissue. Furthermore, it could be combined with the SUSHI labeling technique [42] which, by labeling the extracellular space, enables the investigation of cellular relationships and morphology in living tissue. Our technology can extend the benefits of this labelling technique deep inside the living mouse brain. Altogether, our developments represent the advancement of 3D-STED microscopy into the realm of deep-tissue (*in vivo*) imaging.

## Methods

### Custom-built 3D-2PE-STED microscope for aberration-corrected super-resolution imaging

A detailed schematic of our optical setup is shown in **Fig. 1-figure supplement 1**. Mechanical drawings, parts lists and software are available upon request from the authors.

Our AO-enabled 2PE-STED system features a pulsed 775 nm depletion laser with a pulse length of ∼600-ps (Katana HP, OneFive). The laser beam is expanded to illuminate a spatial light modulator (SLM; X10468-02, Hamamatsu) positioned in a plane conjugate to the objective back aperture. To impart both the vortex and top-hat phase masks on the same depletion beam, we adopt a double-pass SLM configuration as described by Lenz *et al.* [15]. Both SLM phase masks are then imaged onto a 140-actuator deformable mirror (DM; Multi-5.5, Boston Micromachines), a 10 kHz resonance mirror (SC-30, EOPC), a galvanometric (galvo) mirror (6810P, Cambridge Technologies) and finally, the back pupil of a 25×/1.05 NA water immersion objective lens (XLPLM25XWMP2, Olympus), mounted on a custom-built upright microscope stand.

The 2PE beam is supplied by an 80 MHz fs pulsed titanium-sapphire laser (MaiTai HP, Spectra-Physics) that is merged with the STED beam via a dichroic mirror (Dichroic 1; T770dcbpxr, 5 mm thick, Chroma Technology). In the common beam path, the DM corrects for aberrations in the 2PE and STED beams simultaneously, while the resonance and galvo mirrors scan both beams along the fast and slow scanning axes, respectively. Samples were mounted on a *z* piezo stage (P-733.3DD, Physik Instrumente) for scanning in the axial direction.

Fluorescence emission is collected by the same objective that focuses the 2PE and STED beams into the sample. It is split from the depletion and excitation light by a dichroic mirror (Dichroic 3; T650/160dcbpxr, 5 mm thick, Chroma Technology) into two channels. In the image acquisition channel, far-red fluorescence is transmitted by Dichroic 3 and collected by a PMT (H10770PA-40-04, Hamamatsu Corporation) in a non-descanned detection geometry. The PMT is fit with two bandpass emission filters (FF01-624/40, Semrock) to isolate fluorescence emission from the organic dyes ATTO594 and ATTO590. The detected photon counts are then relayed to a pulse discriminator (F-100T, Advanced Research Instruments Corporation) and then to a custom-made circuit board for gated detection (Opsero Electronics). This board is synchronized to the excitation laser trigger signal and it enables software control of the detection window width and delay. This same circuit board is also used to generate a trigger signal for the depletion laser such that its pulses are synchronized with those of the excitation laser. Laser blanking and galvo mirror control during image acquisition is performed by a field-programmable gate array (FPGA) board (PCIe-7841R, National Instruments), while photon counting and image formation is performed by a second FPGA board (PCIe-7852R, National Instruments). Both FPGA boards are synchronized to the motion of the resonance mirror.

In the wavefront sensing channel, fluorescence from the non-linear guide star is reflected by Dichroic 3, descanned, and then coupled out of the common beam path by another dichroic mirror (Dichroic 2; T560lpxrxt, 5 mm thick, Chroma Technology). The fluorescence is then passed through two bandpass emission filters (FF01-510/84, Semrock) and routed to a SHS, which is positioned such that the microlens array (64-483, Edmund Optics) is conjugate to the back focal plane of the objective lens. The SHS spot diagram is acquired by an EMCCD camera (iXon EM+ 897, Andor Technology) positioned at the focal plane of the microlens array and synchronized to image acquisition by the PMT. Though the wavefront sensing channel is spectrally isolated from the non-descanned imaging channel (red emission from ATTO590 or ATTO594), the same 2PE wavelength of 810 nm could be used to excite fluorescence in both channels simultaneously.

### DM and SH Sensor Characterization

The preliminary calibration of the DM was performed prior to integrating the DM into the microscope system. The individual actuator influence functions were measured using an offline Michelson interferometer and decomposed into their constituent Zernike modes as described in [43]. This generated a matrix that, once inverted, could output the appropriate actuator voltage settings for each Zernike mirror mode. The shape of the DM surface could then be controlled by applying any linear combination of these modes.

Once calibrated, the DM was integrated into the microscope system, in a plane conjugate to the objective back pupil. To correct for system aberrations, we imaged 200 nm crimson fluorescent beads (F8806, Invitrogen) embedded in phosphate buffered saline (PBS) containing fluorescein (46955, Sigma-Aldrich) and used a metric-based AO approach to optimize the quality of the bead images [44]. We corrected Zernike mirror modes 4 to 11 using an image brightness metric. Next, we acquired a reference spot diagram by exciting (via 2PE) and descanning the fluorescein emission to the SHS. This reference spot diagram was then used to perform wavefront sensing, as described in [45].

In the final step, the DM was recalibrated *in situ*, using the SHS (rather than an offline interferometer) to decompose the actuator influence functions into Zernike modes. This additional calibration step, performed with the DM in the system, eliminated any discrepancy between the Zernike mode reference frame of the DM (previously defined relative to the interferometer) and that of the SHS.

### Resolution Quantification

The resolution of our system was quantified as described in the main text, using the nested-loop ensemble PSF (NEP) fitting method [33] to account for underlying structure size where appropriate. To quantify the lateral resolution from STEDxy images of microtubules (as shown in Fig. 1(c), left), standard NEP fitting with a Lorentzian-PSF microtubule model function was used. To quantify the lateral and axial resolution from 3D-STED and STEDz image stacks (as shown in Fig. 1(c), middle and right, and Fig. 4(e)), the NEP fitting procedure was extended to quantify the axial PSF size in addition to the lateral PSF and feature sizes.

Multi-axis (lateral and axial) line profiles were extracted such that they intersected at the 3D center of each tubule cross-section. This allowed the algorithm to fit a single tubule diameter shared by both profiles. As a result, the total number of fit parameters that determined the model widths was reduced from 4N to N + 2, where N is the number of line profiles. First, lateral profiles were hand-drawn on an axially-summed or maximum-intensity-projected two-dimensional (2D) image using FIJI. The center positions, angles, and lengths of these profiles were then saved and imported by a Python script. For each manually drawn profile, a preliminary lateral profile was extracted from the axially summed image stack and fit with a Gaussian to determine the lateral coordinates of the tubule center position. Next, a square grid, sized w_*xy*_ × w_*xy*_ (see **Table 1**) with spacings matching the image pixel size (see **Table 2**), was generated about this center position and rotated to match the line orientation. Each slice of the image stack was then interpolated using a bivariate spline of order 3 in both *x* and *y*, sampled using the square grid, and the average value from this interpolation was taken to be the axial profile intensity value at that slice. The extracted axial profile was then fit to a Gaussian to determine the tubule center along *z*. A linear interpolation of the data was sampled over a depth of wz (see **Table 1**) centered on the tubule. This interpolated ministack was then averaged axially to create a 2D image from which the final lateral profile of width w*xy* could be extracted.

**Table 1.** Line Profile Parameters.

| Figure | Imaging Mode | Structure Imaged | Line Profile Widths (nm) | # of FOVs | # of Profiles | PSF FWHM (nm) | Mean Structure Diameter ± Standard Deviation (nm) |
| --- | --- | --- | --- | --- | --- | --- | --- |
| 1(c), left | STED<sub>xy</sub> | Microtubules | $w_{xy} = 1,955^*$<br>$w_z = \text{N/A}$ | 8 | N = 105 | PSF<sub>xy</sub> = 70<br>PSF<sub>z</sub> = N/A | N/A |
| 1(c), middle | 3D-STED | ER tubules | $w_{xy} = 391$<br>$w_z = 400$ | 1 | N = 70 | PSF<sub>xy</sub> = 133<br>PSF<sub>z</sub> = 454 | 107 ± 28 |
| 1(c), middle | 2PE | ER tubules | $w_{xy} = 391$<br>$w_z = 400$ | 1 | N = 70 | PSF<sub>xy</sub> = 354<br>PSF<sub>z</sub> = 1,131 | N/A |
| 1(c), right | STED<sub>z</sub> | ER tubules | $w_{xy} = 391$<br>$w_z = 400$ | 7 | N = 160 | PSF<sub>xy</sub> = 247<br>PSF<sub>z</sub> = 151 | 112 ± 28 |
| 4(e), Day 1 | 3D-STED | Neuron branches | $w_{xy} = 586$<br>$w_z = 600$ | 1 | N = 12 | PSF<sub>xy</sub> = 209<br>PSF<sub>z</sub> = 321 | 366 ± 48 |
| 4(e), Day 3 | 3D-STED | Neuron branches | $w_{xy} = 586$<br>$w_z = 600$ | 1 | N = 7 | PSF<sub>xy</sub> = 160<br>PSF<sub>z</sub> = 320 | 407 ± 47 |

\*average width of hand-drawn line profiles

**Table 2.** Imaging Parameters.

| Figure | Depth<br>($\mu\text{m}$) | Line<br>Accumulations | FOV size<sup>&</sup><br>(pixels)<br>($\mu\text{m}$) | Voxel size<br>(nm) | 810 nm<br>Exc. Power<br>(mW) | 775 nm<br>Depl. Power<br>(mW) |
| --- | --- | --- | --- | --- | --- | --- |
| 1(c)<br>Left | - | 2PE: 100<br>STED: 400 | $1024 \times 1024 \times \text{N/A}$<br>$20 \times 20 \times \text{N/A}$ | $19.5 \times 19.5 \times \text{N/A}$ | 5.20 | Not measured |
| 1(c)<br>Middle | - | 2PE: 300<br>STED: 600 | $512 \times 512 \times 40$<br>$20 \times 20 \times 2$ | $39 \times 39 \times 50$ | 2.30 | Total: 216 |
| 1(c)<br>Right | - | 2PE: 300<br>STED: 200 | $256 \times 256 \times 50$<br>$20 \times 20 \times 2$ | $78 \times 78 \times 40$ | 2.17 | Top-hat: 161 |
| 2(a) | 62 | STED: 200 | $512 \times 512 \times 120$<br>$20 \times 20 \times 6$ | $39 \times 39 \times 50$ | 2PE image:<br>0.85<br>STED image:<br>1.88 | Top-hat: 98 |
| 2(b) | 111 | 2PE: 100<br>STED: 400 | $512 \times 512 \times 80$<br>$20 \times 20 \times 8$ | $39 \times 39 \times 100$ | 3.02 | Not measured |
| 3(a) | 164 | 2PE: 150<br>STED: 500 | $256 \times 256 \times 50$<br>$30 \times 30 \times 10$ | $117 \times 117 \times 200$ | Not measured | Not measured |
| 4(c) | 104 | - | $1024 \times 1024 \times 40$<br>$292 \times 292 \times 40$ | $286 \times 286 \times 1000$ | Not measured | Not measured |
| 4(e) | 76 | See text above | $512 \times 512 \times 80$<br>$20 \times 20 \times 4$ | $39 \times 39 \times 50$ | 2PE image: 1.92<br>STED image: 10.60<br>(Day 3) | Total: 160<br>(Day 3) |

<sup>&</sup>refers to original uncropped image

NEP fitting with a Lorentzian PSF model was used for all STED resolution quantifications. For 2PE resolution quantification, lateral and axial line profiles were extracted as described above, but were fit to a simple Gaussian function. NEP fitting was no longer required since the (close to) diffraction-limited PSF was much larger than the underlying structure size in all three dimensions. In our 2PE data, the simple Gaussian fits resulted in a 30% lower mean mean-squared error than simple Lorentzian fits, indicating that the Gaussian is a better model of our 2PE PSF, as expected.

### Imaging Parameters

Acquisition parameters for all images presented in the main text are summarized in **Table 2**. All images, except for the image in **Fig. 4(c)** and **4(d)**, were acquired using our custom 3D-STED setup using an excitation wavelength of 810 nm and a depletion wavelength of 775 nm. The image in **Fig. 4(c)** was acquired on a commercial 2PE microscope (Ultima Investigator, Bruker Corporation) with multi-color imaging capabilities. An excitation wavelength of 810 nm was also used. **Fig. 4(d)** was acquired on a commercial widefield microscope (DM IL LED, Leica Microsystems), equipped with a CCD camera (DR-328G-C01-SIL Clara, Andor Technology).

The specified imaging depth is measured from the surface of the tissue section or brain to the center of the 3D image stack. “Line Accumulations” refers to the number of times each line in the frame was scanned by the resonance scanner. Fluorescence was only recorded on the forward scan of the mirror. For the image stack in **Fig. 4(e)**, motion artifacts as described in the main text, were prevalent in our initial images. To mitigate this effect, each step in the stack was acquired as a set of 10 individual frames, with the line accumulations set to 20 lines for each frame. Each set of frames was then registered and averaged before being assembled into the final image stack. Excitation and depletion laser powers were measured at the sample, using a power meter (PM100D, Thorlabs) with a photodiode or thermal (S170C or S175C, Thorlabs) sensor.

### Aberration-correction Routine

For all aberration-corrected imaging, spot diagram acquisition was performed prior to image acquisition by scanning the non-linear guide star across a single 2D plane in the middle of the imaged volume. Spot diagrams were analyzed using an algorithm described in [45]. As a minor modification, we localized each spot in the spot diagram using a radial symmetry centers approach [46] rather than centroiding. The calculated phase correction was then applied to the DM for the entire acquisition of the image stack.

The SHS EMCCD exposure times ranged between 1.5 s for the brightest samples and 12 s, if scattering was an issue. However, most spot diagram acquisitions required 5 s exposure times or less. The excitation laser power during spot diagram acquisition were set to be the same as during STED image acquisition, except for the image in **Fig. 2(b)** where the excitation power was increased from 3.02 mW to 11.29 mW for correction.

### Cell Culture and Labelling

An 18 × 18 mm coverslip was sonicated for 15 minutes in potassium hydroxide, rinsed three times with Milli-Q water, sterilized with 100% ethanol, incubated with poly-l-lysine for 10 minutes, and then rinsed once with phosphate-buffered saline (PBS).

For endoplasmic reticulum (ER) labelling, COS7 cells were electroporated with an ER membrane marker (mEmerald-Sec61-C-18, a gift from Michael Davidson, Addgene plasmid #54249) before they were seeded on the coverslip and grown overnight. The cells were fixed with 3% paraformaldehyde (PFA; 15710, Electron Microscopy Science) + 0.1% glutaraldehyde (GA; 16019, Electron Microscopy Science) diluted in PBS that was pre-warmed to 37°C for 15 minutes. The cells were rinsed three times in PBS, permeabilized (PBS + 0.05% IGEPAL CA-630 + 0.05% Trition X-100 + 0.1% Bovine Serum Albumin) for 3 minutes, and then rinsed an additional three times with PBS. The cells were incubated in block buffer (PBS + 0.05% IGEPAL CA-630 + 0.05% TritionX-100+ 5% normal goat serum) for 1 hour then incubated with rabbit anti-GFP antibodies (1:500 dilution in block buffer; A-11122, Invitrogen), overnight at 4°C. The cells were rinsed three times for five minutes each with wash buffer (PBS + 0.05% IGEPAL CA-630 + 0.05% Trition X-100+ 0.2% Bovine Serum Albumin), incubated with secondary anti-rabbit antibodies conjugated to ATTO594 (1:1000 dilution in block buffer; 77671, Sigma-Aldrich) for an hour, and then washed three more times for five minutes each. The cells were post fixed with 3% PFA + 0.1% GA diluted in PBS for 10 minutes, rinsed 3 times with PBS, then mounted were mounted in on a microscope slide using ProLong Diamond Antifade Mountant (P36961, Invitrogen).

For microtubule labelling, COS7 cells were incubated with 0.2% saponin diluted in cytoskeletal buffer (CBS; 10 mM MES pH 6.1, 138 mM NaCl, 3 mM MgCl2, 2 mM EGTA, 320 mM sucrose) that was warmed to 37 °C for 1 minute. The cells were fixed with 3% PFA + 0.1% GA diluted in CBS warmed to 37 °C for 15 minutes, then rinsed 3 times with PBS. Cells were blocked and permeabilized with 3% bovine serum albumin (BSA; 001-000-162, Jackson ImmunoResearch) and 0.2% Triton X-100 (T8787, Sigma-Aldrich) diluted in PBS for 30 minutes. The cells were incubated with mouse anti-α-tubulin antibody (T5168, Sigma-Aldrich) diluted to 1:200 in antibody dilution buffer (1% BSA and 0.2% Triton X-100 in PBS) overnight at 4 °C and then washed three times for five minutes each with wash buffer (0.05% Triton X-100 in PBS). Next, the cells were incubated with anti-mouse secondary antibodies conjugated to ATTO594 (76085-1ML-F, Sigma-Aldrich) for 1 hour and then washed three times for five minutes each with wash buffer. The cells were then post fixed with 3% PFA + 0.1% GA, diluted in CBS for ten minutes and then rinsed three times with PBS. Finally, the cells were mounted in on a microscope slide using ProLong Diamond Antifade Mountant.

### Mouse Skin Tissue Preparation and Labelling

Skin tissue was harvested from a transgenic mouse expressing histone 2B-green fluorescent protein (H2B-GFP) under the control of the keratin 14 (K14) promoter. The tissue was dissected and incubated with 3.5 mg/mL Dispase II (4942078001, Sigma-Aldrich) in PBS for 10 minutes at 37°C. The epidermis was then peeled off of the dermis with fine forceps and fixed in 4% PFA in PBS for 10 minutes. Next, the fixed tissue was incubated in blocking solution (5% normal goat serum, 1% BSA, and 0.2% Triton X-100 in PBS) for 30 minutes, then incubated with anti-GFP nanobodies conjugated to ATTO594 (diluted 1:200 in blocking solution; gba594, ChromoTek) overnight at 4°C. The tissue was then washed in blocking solution 3 times for 10 minutes each, then mounted in ProLong Diamond Antifade Mountant.

For thick tissue preparations, tissue was harvested and dissected as described above, then immediately fixed in 4% PFA in PBS for 1 hour at room temperature. The fixed tissue was then incubated in blocking solution for 6-8 hours at room temperature on a rocking platform. Next, the blocked tissue was incubated with ATTO594-conjugated anti-GFP nanobodies (prepared as described above) for ∼66 hours at room temperature, washed in PBS with 2% Triton X-100 for 6-8 hours, and mounted upside-down (i.e. with the skin surface positioned face-down) in ProLong Diamond Antifade Mountant for imaging.

### Mouse Brain Tissue Preparation and Labelling

A whole mouse brain, fixed in 4% PFA overnight, was sectioned into 300 µm slices using a vibratome (VT1000 S, Leica Biosystems). The sections were then placed in permeabilization buffer (0.5% Triton X-100 in 1× PBS) for 1 hour on a rocking platform at room temperature. Next, the sections were stained with a GFAP polyclonal primary antibody (1:500 dilution; ab7260, Abcam) in antibody dilution buffer (1% BSA + 0.2% Triton X-100 in 1× PBS) first for 24 hours at room temperature on a rocking platform and then for 5 days at 4°C. The sections were subsequently stained with an ATTO594-conjugated secondary antibody (1:500 dilution; 77671, Sigma-Aldrich) for 72 hours at 4°C. After each staining step, the sections were washed 5-7 times with wash buffer (0.05% Triton X-100 in 1× PBS) for 30 minutes each. The samples were then rinsed with PBS and mounted in fluorescein (46955, Sigma-Aldrich) in PBS.

### Animals

All animal procedures described have been approved by the Institutional Animal Care and Use Committee at Yale University. CD1 Mice were purchased from Charles River.

### Plasmid Generation and Viral Vector Production

The AAV-CAG-Halo-GFP viral vector was constructed from the AAV-CAG-GFP plasmid (Addgene plasmid #28014). The GFP sequence was replaced with a Halo-GFP sequence. Recombinant adeno-associated viruses (rAAVs) of serotype 2 were produced and purified following procedures described previously [47], using a two-plasmid helper free system (PlasmidFactory). HEK293T cells were co-transfected with the transfer plasmid, containing the target gene, and the helper plasmid, using JetPrime reagents. Transfected cells were collected 4-5 days later. Virus particles were extracted from the cell lysate and purified by iodixanol gradient ultracentrifugation and titered by transfection assay.

### Neonatal Injection of rAAVs

rAAV viruses were injected into the subarachnoid space of P0 CD1 mice, using a Picospritzer microinjection system (051-0500-900, Parker Hannifin Corporation). Injection pipettes were prepared from glass capillary tubes (3-000-210-G, Drummond Scientific) using a pipette puller (PC-10, Narishige), loaded with the prepared virus solution, and connected to the Picospritzer pipette holder. To prepare for the injection, the mouse pups were placed on ice to induce anesthesia via hypothermia. Next, approximately 1 μl of virus solution was injected into the subarachnoid space of each pup. After the injection, the pups were placed on a warming pad until their body temperature and skin color returned to normal and they began to move. The total number of virus particles injected into a single mouse was approximately 10E7.

### ATTO590 Dye Labelling and Installation of Cranial Window

This procedure was performed at least 21 days after the neonatal injection of rAAVs, and only if the animal recovered normally. The infected mouse was weighed and dosed with buprenorphine (0.1 mg/kg) and ketamine/xylazine (K/X; 100 mg/kg Ketamine, 10 mg/kg Xylazine) via intraperitoneal injection. Furthermore, dexamethasone (2mg/kg) and carprofen (5 mg/kg) were administered via subcutaneous injection. When the mouse was no longer responsive to a firm toe pinch, ophthalmic ointment was applied, the head was shaved and the exposed skin was treated with povidone-iodine solution and cleaned with ethanol. A small piece of skin was then removed to expose the skull, and the membrane tissue covering the skull surface was removed using forceps. The animal was restrained onto a custom stage mount fitted with a heated pad. Next, a dental drill was used to create a 3 mm diameter craniotomy, centered ∼2.5 mm from both Bregma and midline. During this step, the skull was periodically rinsed with sterile PBS to prevent excessive heating. Once the skull was sufficiently thinned, the circular piece of skull was gently lifted and removed using forceps. GELFOAM Sterile Sponge (0315-08, Pfizer Inc.) was used to absorb any bleeding. Next, fine forceps and a needle were used to pierce the dura and gently pull it to one side. 25 µl of ATTO590-CA dye solution was then applied topically onto the exposed brain surface. To prepare the dye solution, 8 µl of 0.6 mM ATTO590-CA solution (molecular weight = 788 g/mol; resuspended in DMSO) was mixed with 40 µl of sterile PBS and 2 µl of Pluronic for a final ATTO590-CA concentration of 96 µM. After incubating for 30 minutes, the area was washed with sterile PBS and the craniotomy was sealed with a piece of #1.5 cover glass, 3 mm in diameter. Two-component silicone putty (Picodent Twinsil, Picodent) was then applied to the circular opening of the head post to protect the area. The animal was then released and allowed to recover in a cage positioned above a heated pad. For animals that underwent multiple imaging sessions over days, buprenorphine was administered twice daily and carprofen (5 mg/kg) was administered once daily for the 2 days following the procedure.

### *in vivo* Imaging

The animal was anesthetized using ketamine/xylazine (100 mg/kg Ketamine, 10 mg/kg Xylazine) via intraperitoneal injection., and when the mouse was no longer responsive to a firm toe pinch, it was restrained onto the custom stage mount via the surgically installed head post. The two-component silicone putty (Picodent Twinsil, Picodent) protecting the circular opening of the head post was removed to expose the glass window, and the window was cleaned with sterile PBS to remove any residue. The custom stage base holding the mouse was then mounted onto the 2PE-STED microscope for *in vivo* super-resolution imaging.

At the end of the imaging session, if the animal was to undergo additional imaging experiments, it was released from the stage mount and allowed to recover in its cage, which was positioned over a heating pad. Otherwise, the mouse was euthanized, either via cardiac perfusion or cervical dislocation.

## Acknowledgements

The authors thank Aarushi Gupta and Dr. Alanna Schepartz for the supply of ATTO590-CA, Dr. Xiang Hao, Dr. Emil Kromann, Dr. Yongdeng Zhang and Mark Lessard for helpful discussions. This work was supported by the National Institutes of Health (1R01NS089734, P30-DK45735, U01 DA047734), the G. Harold & Leila Y. Mathers Charitable Foundation, Yale’s Integrated Graduate Program in Physical and Engineering Biology, the European Research Council AdOMIS (695140), and the Wellcome Trust (203285/B/16/Z, 203285/C/16/Z). J.B. discloses significant financial interest in Bruker Corporation and Hamamatsu Photonics.

## Contributions

M. G. M. V. and J. B. designed the instrument. M. G. M. V. built and optimized the instrument. E. S. A., and M. G. M. V. developed and optimized the microscope control software. M. G. M. V, J. A., M. J. B. and J. B. developed the adaptive optics strategy. M. Z., P. Y., and J. G. optimized the *in vivo* labelling and performed all animal surgeries. D. M. and V. G. prepared the skin tissue samples. O. M. prepared the fixed brain slice samples. P. K. prepared the cultured cell samples. A. E. S. B. developed the NEP fitting code for 3D resolution quantification. M. G. M. V. collected the STED microscopy data. M. G. M. V. and J.B. wrote the manuscript. All authors edited the manuscript.

## Figure Supplements and Movies

**Fig. 1-figure supplement 1.**
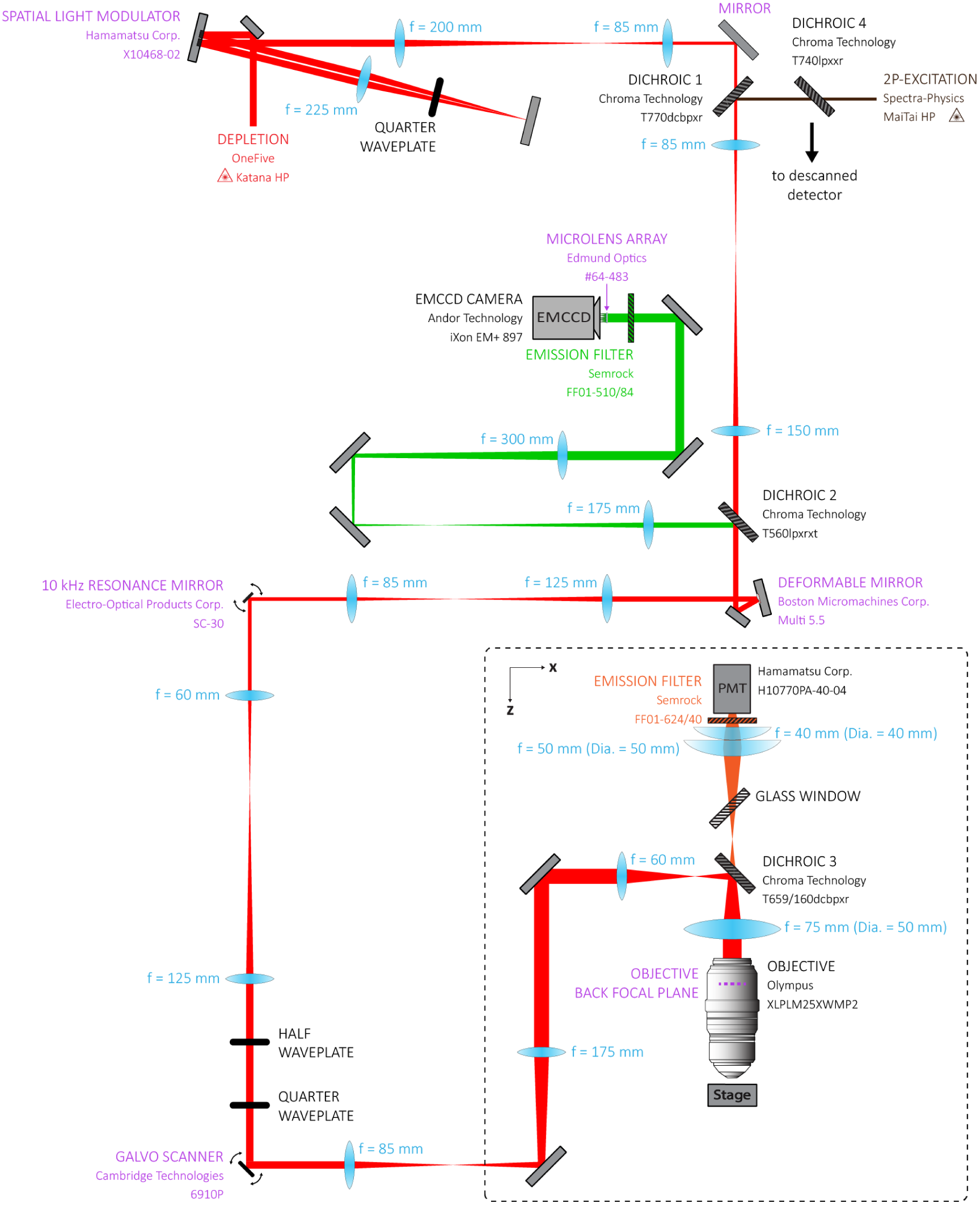
Detailed schematic of our 3D-2PE-STED microscope. See text for details. All optics labeled in purple are positioned in conjugate planes. Dichroic 4 (not discussed in text) is included in the schematic for completeness but was not crucial to the experiments discussed here.

**Supplementary Movie 1. Related to Figure 3. | Aberration-corrected 2PE-STED imaging of ATTO594-labelled astrocytes in fixed mouse brain tissue.**

**Supplementary Movie 2. Related to Figure 4. | Aberration-corrected 2PE-STED imaging of ATTO590-labelled neurons in a living mouse.**

## References

1. S. J. Sahl, S. W. Hell, and S. Jakobs, “Fluorescence nanoscopy in cell biology,” Nat. Rev. Mol. Cell Biol. 18, 685–701 (2017).

2. E. Abbe, “Beiträge zur Theorie des Mikroskops und der mikroskopischen Wahrnehmung,” Archiv für mikroskopische Anatomie 9, 413–418 (1873).

3. L. K. Schroeder, A. E. S. Barentine, H. Merta, S. Schweighofer, Y. Zhang, D. Baddeley, J. Bewersdorf, and S. Bahmanyar, “Dynamic nanoscale morphology of the ER surveyed by STED microscopy,” J. Cell Biol. 218, 83–96 (2018).

4. S. W. Hell, and J. Wichmann, “Breaking the diffraction resolution limit by stimulated emission: stimulated-emission-depletion fluorescence microscopy,” Opt. Lett. 19, 780–782 (1994).

5. S. Dunst, and P. Tomancak, “Imaging flies by fluorescence microscopy: Principles, technologies, and applications,” Genetics 211, 15–34 (2019).

6. B. R. Rankin, G. Moneron, C. A. Wurm, J. C. Nelson, A. Walter, D. Schwarzer, J. Schroeder, D. A. Colon-Ramos, and S. W. Hell, “Nanoscopy in a living multicellular organism expressing GFP,” Biophys. J. 100, L63–L65 (2011).

7. H. Steffens, W. Wegner, and K. I. Willig, “In vivo STED microscopy: A roadmap to nanoscale imaging in the living mouse,” Methods, 10.1016/j.ymeth.2019.1005.1020. (2019).

8. P. Török, and P. R. T. Munro, “The use of Gauss-Laguerre vector beams in STED microscopy,” Opt. Express 12, 3605–3617 (2004).

9. T. A. Klar, and S. W. Hell, “Subdiffraction resolution in far-field fluorescence microscopy,” Opt. Lett. 24, 954–956 (1999).

10. T. J. Gould, D. Burke, J. Bewersdorf, and M. J. Booth, “Adaptive optics enables 3D STED microscopy in aberrating specimens,” Opt. Express 20, 20998–21009 (2012).

11. M. Dyba, and S. W. Hell, “Focal spots of size lambda/23 open up far-field florescence microscopy at 33 nm axial resolution,” Phys. Rev. Lett. 88, 163901 (2002).

12. R. Schmidt, C. A. Wurm, A. Punge, A. Egner, S. Jakobs, and S. W. Hell, “Mitochondrial cristae revealed with focused light,” Nano Lett. 9, 2508–2510 (2009).

13. R. Schmidt, C. A. Wurm, S. Jakobs, J. Engelhardt, A. Egner, and S. W. Hell, “Spherical nanosized focal spot unravels the interior of cells,” Nat. Methods 5, 539–544 (2008).

14. T. A. Klar, S. Jakobs, M. Dyba, A. Egner, and S. W. Hell, “Fluorescence microscopy with diffraction resolution barrier broken by stimulated emission,” Proc. Natl. Acad. Sci. U.S.A. 97, 8206–8210 (2000).

15. M. O. Lenz, H. G. Sinclair, A. Savell, J. H. Clegg, A. C. Brown, D. M. Davis, C. Dunsby, M. A. Neil, and P. M. French, “3-D stimulated emission depletion microscopy with programmable aberration correction,” J. Biophotonics 7, 29–36 (2014).

16. S. Deng, L. Liu, Y. Cheng, R. Li, and Z. Xu, “Effects of primary aberrations on the fluorescence depletion patterns of STED microscopy,” Opt. Express 18, 1657–1666 (2010).

17. J. Heine, C. A. Wurm, J. Keller-Findeisen, A. Schonle, B. Harke, M. Reuss, F. R. Winter, and G. Donnert, “Three dimensional live-cell STED microscopy at increased depth using a water immersion objective,” Rev. Sci. Instrum. 89, 053701 (2018).

18. G. Moneron, and S. W. Hell, “Two-photon excitation STED microscopy,” Opt. Express 17, 14567–14573 (2009).

19. M. J. T. Ter Veer, T. Pfeiffer, and U. V. Nagerl, “Two-photon STED microscopy for nanoscale imaging of neural morphology in vivo,” Methods Mol. Biol. 1663, 45–64 (2017).

20. M. G. Velasco, E. S. Allgeyer, P. Yuan, J. Grutzendler, and J. Bewersdorf, “Absolute two-photon excitation spectra of red and far-red fluorescent probes,” Opt. Lett. 40, 4915–4918 (2015).

21. N. Kilian, A. Goryaynov, M. D. Lessard, G. Hooker, D. Toomre, J. E. Rothman, and J. Bewersdorf, “Assessing photodamage in live-cell STED microscopy,” Nat. Methods 15, 755–756 (2018).

22. D. S. Richardson, and J. W. Lichtman, “Clarifying tissue clearing,” Cell 162, 246–257 (2015).

23. M. Schwertner, M. Booth, and T. Wilson, “Characterizing specimen induced aberrations for high NA adaptive optical microscopy,” Opt. Express 12, 6540–6552 (2004).

24. M. Schwertner, M. J. Booth, M. A. A. Neil, and T. Wilson, “Measurement of specimen-induced aberrations of biological samples using phase stepping interferometry,” J. Microsc. 213, 11–19 (2004).

25. B. R. Patton, D. Burke, D. Owald, T. J. Gould, J. Bewersdorf, and M. J. Booth, “Threedimensional STED microscopy of aberrating tissue using dual adaptive optics,” Opt. Express 24, 8862–8876 (2016).

26. P. Zdankowski, D. McGloin, and J. R. Swedlow, “Full volume super-resolution imaging of thick mitotic spindle using 3D AO STED microscope,” Biomed. Opt. Express 10, 1999–2009 (2019).

27. R. Aviles-Espinosa, J. Andilla, R. Porcar-Guezenec, O. E. Olarte, M. Nieto, X. Levecq, D. Artigas, and P. Loza-Alvarez, “Measurement and correction of in vivo sample aberrations employing a nonlinear guide-star in two-photon excited fluorescence microscopy,” Biomed. Opt. Express 2, 3135–3149 (2011).

28. K. Wang, D. E. Milkie, A. Saxena, P. Engerer, T. Misgeld, M. E. Bronner, J. Mumm, and E. Betzig, “Rapid adaptive optical recovery of optimal resolution over large volumes,” Nat. Methods 11, 625–628 (2014).

29. W. Zheng, Y. Wu, P. Winter, R. Fischer, D. D. Nogare, A. Hong, C. McCormick, R. Christensen, W. P. Dempsey, D. B. Arnold, J. Zimmerberg, A. Chitnis, J. Sellers, C. Waterman, and H. Shroff, “Adaptive optics improves multiphoton super-resolution imaging,” Nat. Methods 14, 869–872 (2017).

30. R. Turcotte, Y. Liang, M. Tanimoto, Q. Zhang, Z. Li, M. Koyama, E. Betzig, and N. Ji, “Dynamic super-resolution structured illumination imaging in the living brain,” Proc. Natl. Acad. Sci. U.S.A. 116, 9586–9591 (2019).

31. K. Wang, W. Sun, C. T. Richie, B. K. Harvey, E. Betzig, and N. Ji, “Direct wavefront sensing for high-resolution in vivo imaging in scattering tissue,” Nat. Commun. 6, 7276 (2015).

32. F. Bottanelli, E. B. Kromann, E. S. Allgeyer, R. S. Erdmann, S. Wood Baguley, G. Sirinakis, A. Schepartz, D. Baddeley, D. K. Toomre, J. E. Rothman, and J. Bewersdorf, “Two-colour live-cell nanoscale imaging of intracellular targets,” Nat. Commun. 7, 10778 (2016).

33. A. E. S. Barentine, L. K. Schroeder, M. Graff, D. Baddeley, and J. Bewersdorf, “Simultaneously measuring image features and resolution in live-cell STED images,” Biophys. J. 115, 951–956 (2018).

34. W. R. Zipfel, R. M. Williams, and W. W. Webb, “Nonlinear magic: multiphoton microscopy in the biosciences,” Nat. Biotechnol. 21, 1369 (2003).

35. R. J. Noll, “Zernike polynomials and atmospheric-turbulence,” J. Opt. Soc. Am. 66, 207–211 (1976).

36. J. M. Masch, H. Steffens, J. Fischer, J. Engelhardt, J. Hubrich, J. Keller-Findeisen, E. D’Este, N. T. Urban, S. G. N. Grant, S. J. Sahl, D. Kamin, and S. W. Hell, “Robust nanoscopy of a synaptic protein in living mice by organic-fluorophore labeling,” Proc. Natl. Acad. Sci. U.S.A. 115, E8047–E8056 (2018).

37. G. V. Los, L. P. Encell, M. G. McDougall, D. D. Hartzell, N. Karassina, C. Zimprich, M. G. Wood, R. Learish, R. F. Ohana, M. Urh, D. Simpson, J. Mendez, K. Zimmerman, P. Otto, G. Vidugiris, J. Zhu, A. Darzins, D. H. Klaubert, R. F. Bulleit, and K. V. Wood, “HaloTag: a novel protein labeling technology for cell imaging and protein analysis,” ACS Chem. Biol. 3, 373–382 (2008).

38. R. Turcotte, Y. Liang, and N. Ji, “Adaptive optical versus spherical aberration corrections for in vivo brain imaging,” Biomed. Opt. Express 8, 3891–3902 (2017).

39. G. L. Galiñanes, P. J. Marchand, R. Turcotte, S. Pellat, N. Ji, and D. Huber, “Optical alignment device for two-photon microscopy,” Biomed. Opt. Express 9, 3624–3639 (2018).

40. G. Lukinavicius, K. Umezawa, N. Olivier, A. Honigmann, G. Yang, T. Plass, V. Mueller, L. Reymond, I. R. Correa, Jr., Z. G. Luo, C. Schultz, E. A. Lemke, P. Heppenstall, C. Eggeling, S. Manley, and K. Johnsson, “A near-infrared fluorophore for live-cell super-resolution microscopy of cellular proteins,” Nat. Chem. 5, 132–139 (2013).

41. J. B. Grimm, A. K. Muthusamy, Y. Liang, T. A. Brown, W. C. Lemon, R. Patel, R. Lu, J. J. Macklin, P. J. Keller, N. Ji, and L. D. Lavis, “A general method to fine-tune fluorophores for live-cell and in vivo imaging,” Nat. Methods 14, 987–994 (2017).

42. J. Tonnesen, V. Inavalli, and U. V. Nagerl, “Super-Resolution Imaging of the Extracellular Space in Living Brain Tissue,” Cell 172, 1108–1121 e1115 (2018).

43. J. Antonello, T. van Werkhoven, M. Verhaegen, H. H. Truong, C. U. Keller, and H. C. Gerritsen, “Optimization-based wavefront sensorless adaptive optics for multiphoton microscopy,” J. Opt. Soc. Am. A 31, 1337–1347 (2014).

44. D. Debarre, E. J. Botcherby, T. Watanabe, S. Srinivas, M. J. Booth, and T. Wilson, “Image-based adaptive optics for two-photon microscopy,” Opt. Lett. 34, 2495–2497 (2009).

45. J. Antonello, “Optimisation-based wavefront sensorless adaptive optics for microscopy,” PhD Dissertation, Delft University of Technology (2014).

46. R. Parthasarathy, “Rapid, accurate particle tracking by calculation of radial symmetry centers,” Nat. Methods 9, 724–726 (2012).

47. D. Grimm, M. A. Kay, and J. A. Kleinschmidt, “Helper virus-free, optically controllable, and two-plasmid-based production of adeno-associated virus vectors of serotypes 1 to 6,” Mol. Ther. 7, 839–850 (2003).

